# Hepatitis C virus NS5A is not an effective antagonist of the canonical PKR-eIF2α pathway

**DOI:** 10.64898/2026.10.07.757322

**Authors:** Carla E. Gallardo Flores, Lindsay Yu, Elana R. Kertzman, Paul Rothhaar, Volker Lohmann, Che C. Colpitts

## Abstract

Protein kinase R (PKR) is an antiviral sensor that induces translational shutdown and stress granule (SG) formation through the phosphorylation of eIF2α. Although previous studies have proposed that hepatitis C virus (HCV) non-structural protein 5A (NS5A) inhibits PKR dimerization, other studies consistently observe PKR activation and SG formation during HCV infection. To address this discrepancy, we interrogated the ability of NS5A from different genotypes and strains to regulate canonical PKR-eIF2α activation in a variety of cell culture models. Regardless of the genotype or strain tested, HCV NS5A failed to inhibit PKR dimerization or dsRNA-induced PKR activation, nor did it rescue downstream steps in the PKR-eIF2α pathway, such as translation halt and SG formation, in contrast to known viral PKR antagonists. Similarly, NS5A did not antagonise PKR activation when expressed in the context of other HCV non-structural proteins. We also tested the role of the host factor cyclophilin A (CypA), which interacts with both NS5A and PKR. However, the absence of CypA did not affect downstream PKR signaling or the activity of viral PKR antagonists or NS5A. Collectively, these findings indicate that NS5A does not effectively antagonise the PKR-eIF2α pathway and further supports previous findings that HCV replication occurs in the presence of an active PKR response.

## INTRODUCTION

An estimated 50 million people worldwide are living with chronic hepatitis C virus (HCV) infection, with approximately 1 million new infections or reinfections occurring each year [1]. Thus, the prevalence of chronic HCV infection remains high, despite the availability of effective direct-acting antivirals (DAAs). Chronic HCV infection increases the risk for hepatocellular carcinoma (HCC), the third highest cause of cancer-related mortality, and although DAAs are highly effective at curing HCV, liver damage is not reversed, and HCC risk can persist [2].

HCV is an enveloped virus with a positive-sense single-stranded RNA (+ssRNA) genome that encodes a polyprotein that is processed into three structural and seven non-structural proteins [3, 4]. Following entry and uncoating, the +ssRNA is directly recognized by the cellular ribosomes through an internal ribosome entry site (IRES) in the 5′ untranslated region of the viral genome, which bypasses the need for the 5′ 7meG-cap structure and produces a single polyprotein precursor of ∼3000 amino acids [5]. Finally, this polyprotein is cleaved by viral and cellular proteases into the structural and non-structural proteins that mediate viral RNA replication and particle assembly [4]. Of these proteins, non-structural protein 5A (NS5A) has key roles in host modulation.

HCV is classified in seven genotypes and several subtypes. Historically, HCV genotype 1 subtype b (HCV-gt1b) and genotype 3 have been associated with a higher risk of developing HCC in some patient cohorts [6, 7]. However, the mechanisms underlying genotypic differences in HCC risk are not fully understood. Previous studies have demonstrated that patients with HCV-related HCC present increased levels of protein kinase R (PKR) in malignant and surrounding non-malignant tissues [8]. Modulation of endogenous PKR can also affect HCC progression *in vitro,* and PKR inhibition in mouse xenograft models leads to a reduction in tumor size and longer survival of mice, indicating a relationship between PKR regulation and HCC progression [9, 10].

PKR is a serine/threonine kinase that, upon sensing double-stranded RNA (dsRNA), dimerizes and becomes autophosphorylated [11]. Activated PKR phosphorylates and inhibits the eukaryotic translation initiation factor 2α (eIF2-α), causing a halt in translation [12]. Subsequently, halted translation complexes aggregate via liquid-liquid phase separation to give rise to the formation of structures termed stress granules (SGs) [13]. PKR is also implicated in activation of the antiviral transcription factor interferon regulatory factor 1 (IRF1) [14], as well as nuclear factor kappa-light-chain-enhancer of activated B cells (NF-κB) [15–17]. Given the key role of PKR in sensing and restricting virus infection, many unrelated viruses have evolved different mechanisms to evade or inhibit PKR [12]. For example, influenza A virus (IAV) non-structural protein 1 (NS1), poxvirus K3L, and Ebola virus VP35 inhibit PKR activation [18–20]. However, in the case of HCV infection, the literature regarding the role of PKR is conflicting.

Early studies reported that HCV NS5A from genotype 1b binds and inhibits PKR *in vitro* by disrupting PKR dimerization [21, 22]. The proposed PKR binding domain contains the interferon (IFN)-sensitivity determining region (ISDR) and can be found in low complexity sequence I and domain II of NS5A, which historically was associated with responsiveness to IFN therapy in some patient groups [23], and recently was found to have a central role in regulating viral replication fitness [24]. Nevertheless, infection of cells with cell culture-derived infectious HCV (HCVcc) of the genotype 2a JFH1 strain leads to robust PKR activation and PKR-dependent SG formation, which is enhanced by interferon (IFN) [25–27]. In this context, PKR activation may be proviral. For example, SG-associated proteins such as eukaryotic initiation factor 3 subunit η (eIF3η) and Ras-GTPase-activating protein-binding protein 1 (G3BP1) promote HCV infection by regulating replication, assembly and egress [27, 28]. Oscillation of SG assembly and disassembly in HCV-infected cells is thought to promote cell survival and enable persistent HCV infection [26]. Furthermore, PKR activation hampers the innate immune response by inhibiting cap-dependent translation of interferon (IFN) and IFN-stimulated genes (ISGs) [25, 29], without impairing IRES-driven viral protein synthesis. Overall, a clear understanding of the role of NS5A in regulating PKR during HCV infection is lacking, and potential genotypic differences remain to be investigated.

The proposed PKR binding domain within NS5A is near the binding site for cyclophilin A (CypA) [30], a host peptidylprolyl isomerase that is a well-established host factor for HCV infection [31, 32]. We previously showed that cyclophilin inhibitors, including cyclosporine A (CsA), inhibit HCV replication and infection less potently in cells lacking PKR [33], leading us to hypothesize that disruption of the CypA-NS5A interaction by CsA may rescue PKR function to further restrict HCV infection. In this manuscript, we sought to characterize how NS5A from different HCV genotypes and strains affects PKR activation and canonical downstream translational halt to inform our understanding of the potential role of CypA. However, we found that NS5A, regardless of the tested genotypes and strains, does not effectively inhibit canonical PKR activation or downstream translational repression and stress granule formation. Overall, our data do not support a major role for NS5A as a direct antagonist of classical PKR-eIF2α-mediated translational halt in cell culture models.

## METHODS

### Cell lines

Huh7.5.1 (APC167) cells were obtained from Apath LLC. Huh7-Lunet/T7 cells stably expressing the T7 RNA polymerase were described previously [34]. HEK293T/17 (CRL-11268) cells were obtained from American Type Culture Collection. All cell lines were cultured in complete Dulbecco’s Modified Eagle Medium (DMEM) (Thermo Scientific, 11995065) containing 10% fetal bovine serum (FBS) (Corning, 35-077-CV) and 1% penicillin/streptomycin (Thermo Scientific, 15140122). All cells were cultured at 37 °C in the presence of 5% CO_2_.

### Drugs and reagents

Coumermycin A1 was obtained from Cedarlane (HY-N7452-1MG) and used at 100 ng/mL unless otherwise stated. Dimethyl sulfoxide (DMSO) (Thermo Scientific, D12345) served as the vehicle control. Poly(I:C) was obtained from Invivogen (tlrl-pic) and used at 2 µg/mL.

### Plasmids and cloning

All plasmids were propagated in Stabl competent *Escherichia coli* (NEB, C3040H). The thymidine kinase (TK) promoter-driven Renilla luciferase (TK-RLuc) reporter plasmid has been described previously [35]. The pCMV-Gaussia luciferase (pCMV-GLuc) reporter plasmid was kindly provided by Dr. Craig McCormick (Dalhousie University, Canada). The pSG5-GyrB-PKR-WT and pSG5-GyrB-PKR-K296H plasmids were kindly shared by Dr. Thomas Dever (National Institutes of Health, United States) [36]. pcDNA3.1 plasmids encoding C-terminally FLAG-tagged NS5A from HCV strains H77, GLT1, J4 and JFH1 were generated by HiFi assembly (New England Biolabs (NEB), E2621S) using primers designed by NEBuilder and viral genome plasmids expressing either full-length GLT1 [37] or J6/JFH with chimeric NS5A sequences [38] as the PCR template. Mammalian expression plasmids encoding sheeppox virus (SPPV) K3L and influenza A/Puerto Rico/08/1934 (PR8) NS1 were synthesized by Genscript. The pTM vectors encoding HCV NS3-3′ from different genotypes were previously described [39]. We previously described the pLentiCRISPRv2 plasmid encoding guide RNA targeting CypA [33], which was used alongside psPAX2 packaging (Addgene #12260) and pCMV-VSV-G envelope (Addgene # 8454) plasmids to produce lentivirus.

### Generation of stable cell lines by lentiviral transduction

Lentiviral particles were generated and used to transduce HEK293T/17 cells as described previously [35]. Transduced cells were selected in the presence of 1 μg/mL puromycin and clonal cell lines were generated by limiting dilution in 96-well plates.

### HCV propagation and infection

The J6/JFH1 plasmid was acquired from Apath LLC. The plasmid was linearized by digestion with XbaI and purified with the PCR and DNA cleanup kit (NEB, T1030S). Next, viral RNA was obtained by *in vitro* transcription (NEB, E2040S), and RNA was purified (NEB, T2050S), aliquoted, and stored at -80°C.

Huh7.5.1 cells were seeded in 6-well plates at 4 x 10^5^ cells/well. The next day, cells were transfected with 2.5 μg/well of *in vitro* transcribed HCVcc RNA using Lipofectamine 2000 (Invitrogen, 11668019) according to the manufacturer’s instructions. Cells were incubated at 37°C for 4 hours and then media was replaced with DMEM containing 2% of FBS. After 72 h, transfected cells were transferred to a 100-mm round dish and incubated with DMEM containing 5% FBS at 37°C. After 96 h, supernatant containing HCVcc virus was centrifuged to eliminate cellular debris and passed through a 0.45 µm filter, then aliquoted and stored at -80°C.

For large-scale virus propagation, 3 x 10^5^ Huh7.5.1 cells were seeded in a 6-well plate. The next day, cells were infected with HCVcc (MOI ∼0.003) diluted in serum-free media. After 4 h, the inoculum was removed, and media was replaced with complete DMEM. After 72 h, cells were split and expanded into larger flasks, and after another 72 h, the process was repeated. Supernatant was collected and centrifuged to remove cellular debris and passed through a 0.45 µm filter. Finally, the virus was aliquoted and stored at -80°C.

### HCVcc titration by focus forming assay

Huh7.5.1 cells were seeded at a density of 1 x 10^5^ cells/well in 24-well plates. The next day, HCVcc-containing supernatant was serially diluted in serum-free DMEM. Wells were washed once with PBS and infected using 150 µL/well diluted virus at 37°C. After 4 h, the inoculum was removed, and DMEM containing 5% FBS was added. After 72 h of incubation, infected cells were fixed with 10% formalin for 1 hour. Next, cells were blocked and permeabilized with blocking solution (PBS containing 5% FBS and 0.1% Triton-X100) for 1 h. Cells were then incubated in the presence of a primary antibody against HCV core protein at 4°C overnight. Subsequently, cells were washed and incubated with secondary antibody for 45 minutes. Finally, foci of infected cells were visualized and counted under a Nikon Eclipse Ts2-FL inverted microscope.

### Antibodies

For western blot, primary antibodies targeting HCV core (Anogen MO-I40015B, dilution 1:1000), HCV NS5A (Sigma-Aldrich MABF2775, dilution 1:1,000), α-tubulin (Invitrogen MA531466, dilution 1:10,000), β-actin (Abcam ab8226, dilution 1:10,000), cyclophilin A (Cell Signaling Technology (CST) 2175S, dilution 1:1000), PKR (Abcam ab32052, dilution 1:5,000), phospho-PKR T446 (Abcam ab32036, dilution 1:1,000), eIF2α (CST 5324S, dilution 1:5,000), and phospho-eIF2α (CST 3398S, dilution 1:1,000) were used. The secondary antibodies used were IRDye-680 (red) goat anti-mouse (LICOR 926-68070, dilution 1:10,000) and IRDye-800 (green) (LICOR 926-32211, dilution 1:10,000). For immunofluorescence, the primary antibodies used were HCV core (Anogen MO-I40015B, dilution 1:200), HCV NS5A (Sigma-Aldrich MABF2775, dilution 1:100), eIF3η (Santa Cruz Biotechnology SC-137214), and G3BP1 (CST E9G1M, dilution 1:500). Secondary antibodies Alexa Fluor 488 anti-mouse (CST 4408S, dilution 1:1000) and Alexa Fluor 594 anti-rabbit (CST 8889S, dilution 1:1000) were also used.

### Western blot

For western blot sample collection, cells were scraped into 100 µL of ice-cold PBS containing protease (Roche, 11836170001) and phosphatase (Roche, 4906845001) inhibitors and transferred to microcentrifuge tubes containing 30 µL of 4X SDS-PAGE loading buffer (200 mM Tris pH 6.8, 8% SDS, 0.4% bromophenol blue, 40% glycerol, and 2% β-mercaptoethanol). Next, samples were heated at 95°C for 5 minutes and separated by SDS-PAGE. Subsequently, samples were transferred to nitrocellulose membranes using the TransBlot Turbo System (Bio-Rad) according to manufacturer instructions (25 V, 1.0 A for 30 min). Membranes were blocked in 1X TBS-T (Tris-buffered saline with 0.1% Tween-20) containing 3% BSA for 1 h and subsequently incubated with primary antibodies diluted in TBS-T 3% BSA solution at 4°C overnight. Membranes were washed and then incubated for 1 h at room temperature with Licor IRDye-680 (red) goat anti-mouse and IRDye-800 (green) goat anti-rabbit secondary antibodies diluted 1:10,000. Finally, the membranes were washed and visualized using a Licor Odyssey Clx imager.

### Immunofluorescence

Cells originally seeded onto glass coverslips were fixed using 10% of formalin solution, washed three times with PBS, and blocked and permeabilized using blocking solution for 1 h. Subsequently, cells were incubated with primary antibodies diluted in blocking solution. After overnight incubation at 4°C, cells were washed three times with PBS, incubated with secondary antibodies diluted in blocking solution for 45 minutes, and washed again. Finally, glass coverslips were mounted on microscope slides using FluoroMount G with DAPI (Invitrogen 501128966) and visualized at 63× magnification under water immersion using a Leica Mica confocal microscope.

### Luciferase reporter assays

For protein translation luciferase reporter assays, 500 ng of pcDNA plasmids or GyrB-PKR constructs were transfected together with 100 ng of pCMV-GLuc or 10 ng of TK-RLuc, using FuGENE-6 Transfection Reagent (Promega E2692) in HEK293T/17 cells or TransIT-LT1 (MirusBio MIR-2304) in Huh7-derived cell lines. At the end of the experiment, for TK-Rluc, cells were lysed with 200 μL/well of 1X passive lysis buffer (Promega E1941) or 1X universal lysis buffer (NanoLight #333), and for pCMV-GLuc, cell supernatant was collected. Next, 30 μL of cell lysate or supernatant was transferred into a 96-well white plate, and Renilla or Gaussia luciferase activity was measured by injecting 50 μL of prepared 1 mg/mL coelenterazine substrate (NanoLight #303) into each well using a GloMax plate reader.

### Statistical analysis

All statistical analysis was performed using GraphPad Prism for Mac version 10.1.1. Statistical significance was assessed by two-way ANOVA with Sidak’s multiple comparisons post-hoc test (*p<0.05, **p<0.01, ***p<0.0005, ****p<0.0001).

## RESULTS

### PKR is activated in HCV JFH1-infected cells

While it has been reported that HCV NS5A antagonizes PKR activation [21, 22], PKR is activated during HCVcc infection even with robust NS5A expression [25, 40]. To confirm these observations, we infected Huh7.5.1 cells with HCVcc (J6/JFH1) at a multiplicity of infection (MOI) of 0.1, 0.5, or 1 focus-forming unit (ffu) per cell. At 72 h post-infection, PKR activation was assessed by western blot analysis using an antibody specific for phosphorylated PKR (P-PKR). PKR phosphorylation was increased in HCV-infected cells, indicating activation of PKR during HCV infection **(Figure 1A)**. Since PKR activation is associated with stress granule (SG) assembly, we also examined SG formation in infected cells. SGs were readily detected in HCV-infected cells **(Figure 1B)**, consistent with previous literature [27].

**Figure 1.**
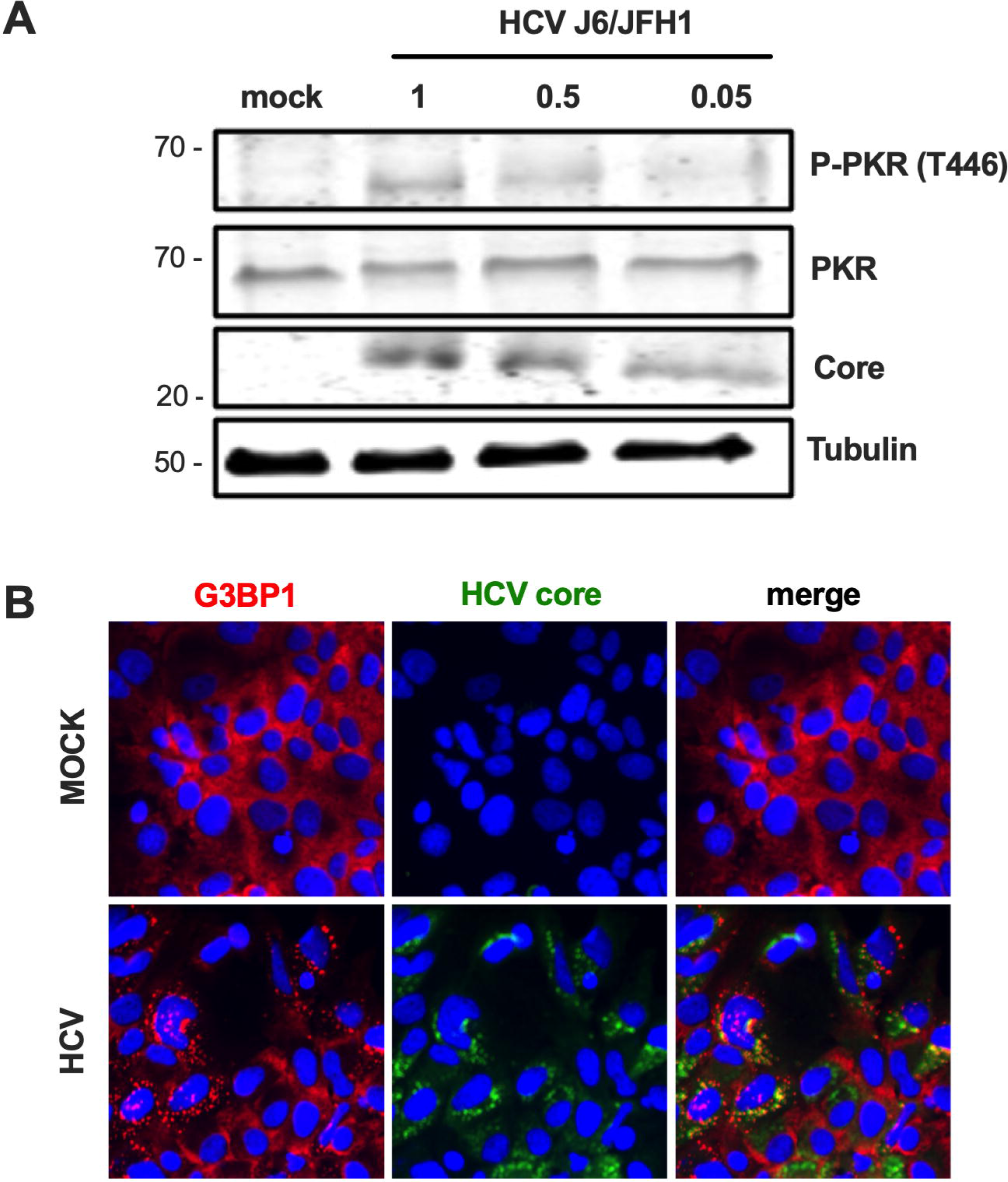
HCV infection leads to PKR activation and SG formation. Huh7.5.1 cells were infected with HCVcc J6/JFH1 at an MOI of 1, 0.5, and 0.05 ffu/mL. **(A)** PKR activation was assessed at 72 hours post-infection by western blot using antibodies for PKR (P-PKR) phosphorylated at position T446, total PKR, HCV core protein, and tubulin. **(B)** SG formation in infected cells was determined by immunofluorescence using antibodies against G3BP1 (red) and HCV core protein (green), with DAPI nuclear stain.

### NS5A does not inhibit selective GyrB-PKR activation

Early studies using yeast models and bacteriophage λ-based genetic assays in *E. coli* proposed that NS5A from IFN-resistant genotype 1b strains binds the catalytic domain of PKR independently of dsRNA and reduces its dimerization [22]. Therefore, to specifically assess the effect of NS5A from different genotypes on PKR dimerization, we employed a selective PKR dimerization construct. In this construct, the dsRNA-binding domain of wild-type PKR (GyrB-PKR-WT) or the kinase-dead mutant PKR-K296H (GyrB-PKR-K296H) is replaced with the *Escherichia coli* Gyrase B (GyrB) domain [36]. Addition of coumermycin A1 induces GyrB-mediated dimerization, thereby triggering PKR autophosphorylation and activation in the absence of dsRNA **(Figure 2A)**.

**Figure 2.**
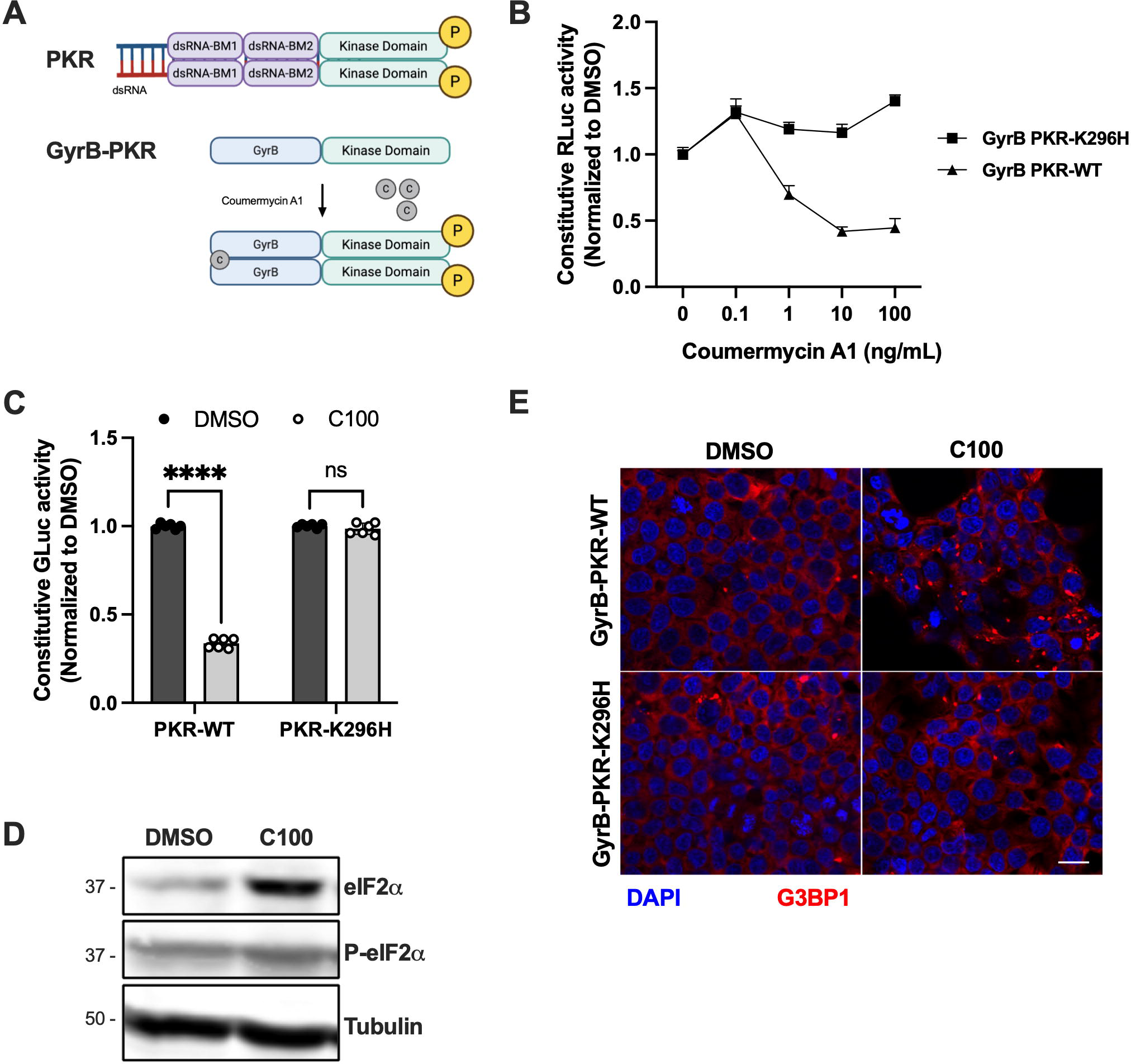
Selective activation of GyrB-PKR. **(A)** Schematic representation of GyrB-PKR construct, where the dsRNA-binding domain of PKR is replaced with GyraseB, allowing PKR dimerization to be induced by coumermycin A1 treatment, independently of dsRNA. **(B-C)** HEK293T/17 cells were co-transfected with constitutive TK-RLuc (B) or pCMV-GLuc (C) reporter plasmid and GyrB-PKR wild type or kinase dead constructs (GyrB-PKR-WT or GyrB-PKR-K296H, respectively). After 24 hours, cells were treated with the indicated concentrations of coumermycin A1. The next day, luciferase reporter activity was measured as a proxy for protein synthesis. **(D-E)** HEK293T-17 cells were co-transfected with GyrB-PKR-WT or GyrB-PKR-K296H 24 h prior to treatment with coumermycin A1 (100 ng/mL). After 2 hours of treatment, cells were processed for western blot to detect EIF2α phosphorylation **(D)**. After 24 hours of treatment, SG formation was assessed by immunofluorescence using antibodies for G3BP1 (red) with DAPI nuclear stain **(E)**. A representative blot and image are shown. Graphs show means +/- standard deviation from at least two independent experiments performed in technical triplicate.

To validate this system, we transfected HEK293T/17 cells with either GyrB-PKR-WT or GyrB-PKR-K296H together with constitutive thymidine kinase promoter-driven Renilla luciferase (TK-RLuc) or constitutive CMV promoter-driven Gaussia luciferase (pCMV-GLuc) reporter plasmids. The following day, cells were treated with increasing concentrations of coumermycin A1 for an additional 24 h. Coumermycin A1 treatment caused a dose-dependent reduction in constitutive luciferase activity, reflecting decreased protein synthesis, in cells expressing GyrB-PKR-WT, whereas no significant reduction was observed in cells expressing the kinase-dead GyrB-PKR-K296H mutant **(Figure 2B-C)**, consistent with PKR-dependent inhibition of protein translation. To further confirm PKR activation, we examined phosphorylation of eIF2α, a canonical downstream target of PKR. Increased eIF2α phosphorylation was detected in cells expressing GyrB-PKR-WT following coumermycin A1 treatment compared to DMSO-treated controls **(Figure 2D)**. In addition, immunostaining for the stress granule marker G3BP1 revealed increased SG formation in cells expressing GyrB-PKR-WT relative to GyrB-PKR-K296H in the presence of coumermycin A1 **(Figure 2E)**. Collectively, these results validate the chemically inducible PKR dimerization system as a robust model to study downstream PKR signaling, supporting its use for subsequent analyses of NS5A-mediated regulation of PKR.

Many viruses have evolved antagonists of PKR. Influenza A virus NS1 (IAV-NS1) inhibits PKR activation upstream of dimerization by binding to the N-terminal region of PKR to block its activation by dsRNA or the protein activator of PKR (PACT) [18, 41]. Poxvirus K3L (Pox-K3L) functions as a pseudo-substrate that competitively inhibits phosphorylation of the PKR substrate eIF2α [19]. The sheeppox virus (SPPV) K3L protein is a particularly effective antagonist of human PKR [42]. We therefore used these well-characterized PKR antagonists as controls to help determine whether and how HCV NS5A inhibits PKR. We focused on NS5A from representative HCV strains, including H77 (genotype 1a), J4 (genotype 1b) and JFH1 (genotype 2a), as well as GLT1, a recently described genotype 1b clinical isolate that has accumulated mutations in the ISDR conferring enhanced replication fitness [24, 37].

HEK293T/17 cells were co-transfected with pcDNA expression plasmids encoding GFP control, SPPV-K3L, IAV-NS1 or HCV-NS5A from the indicated strains together with GyrB-PKR-WT and the constitutive pCMV-GLuc reporter plasmid. Twenty-four hours later, cells were treated with either DMSO or coumermycin A1 for an additional 24 h. As expected, expression of SPPV-

K3L rescued luciferase activity following coumermycin A1 treatment, consistent with its ability to inhibit PKR signaling downstream of the dimerization step **(Figure 3A)**. In contrast, HCV NS5A did not restore luciferase expression relative to the GFP control, regardless of genotype **(Figure 3A)**. NS5A expression was confirmed by western blot **(Figure 3B)**. Similarly, IAV-NS1 failed to rescue translation in the context of chemically induced PKR dimerization, reflecting its established role in inhibiting PKR activation upstream of the dimerization step **(Figure 3A).**

**Figure 3.**
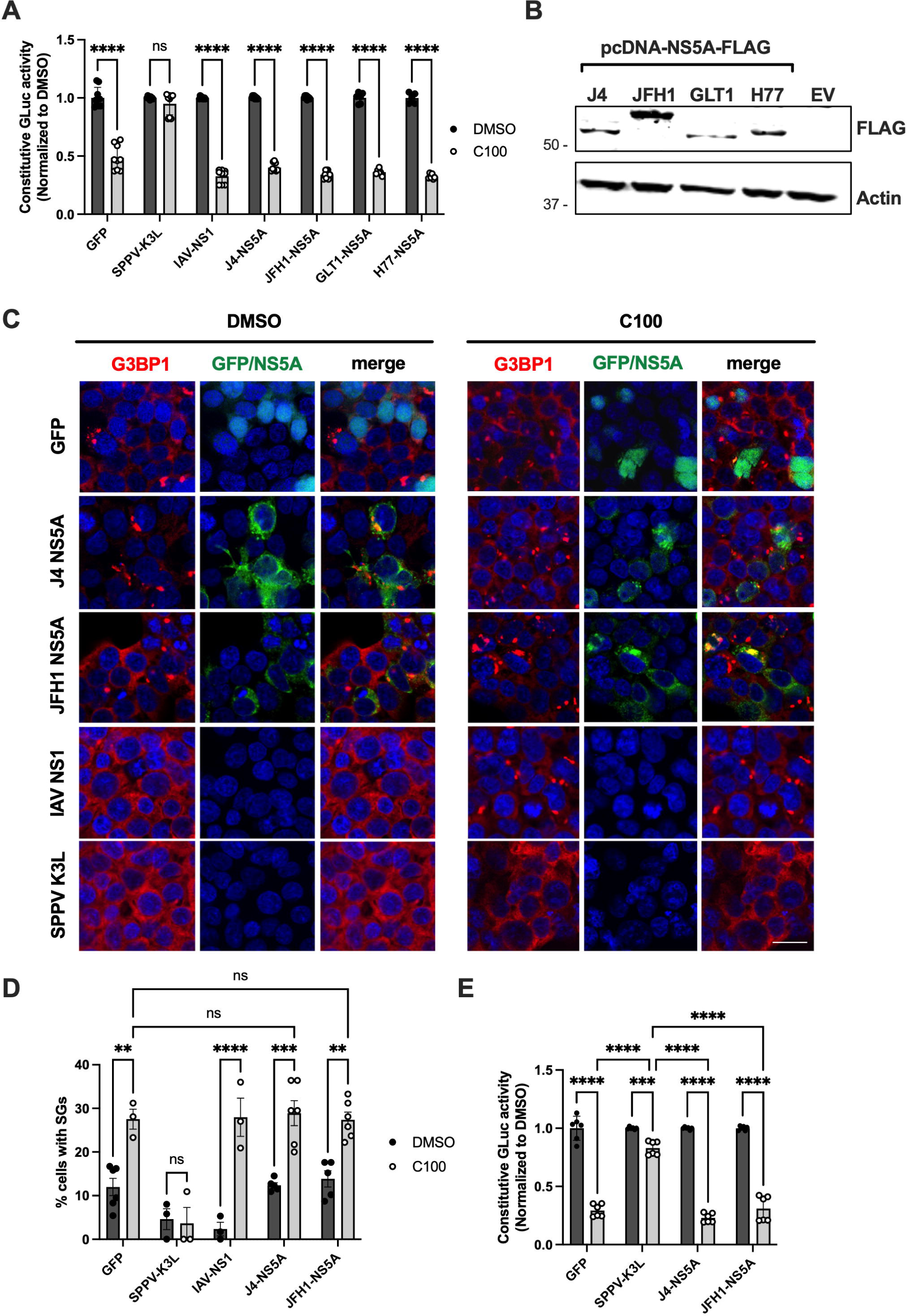
NS5A does not inhibit PKR dimerization or downstream signalling. **(A-D)** HEK293T/17 cells were co-transfected with 500 ng of pcDNA expression plasmids encoding GFP, SPPV-K3L, IAV-NS1, JFH1-NS5A-FLAG, J4-NS5A-FLAG, GLT1-NS5A-FLAG or H77-NS5A-FLAG, together with 500 ng of GyrB-PKR-WT **(A-D)** and the constitutive pCMV-Gluc reporter plasmid **(A-B)**. The next day, cells were treated with DMSO vehicle or coumermycin A1 (100 ng/mL) for an additional 24 h and luciferase activity was measured and normalized to DMSO-treated controls **(A)**. Expression of NS5A-FLAG from different genotypes was confirmed by western blot **(B)**. **(C-D)** SG formation was assessed in cells expressing GFP, J4-NS5A-FLAG, JFH1-NS5A-FLAG, IAV-NS1 or SPPV K3L by immunofluorescence for G3BP1 (red) **(C)**. **(D)** The percentage of cells with SGs was quantified in at least three independent fields of view from two independent experiments. **(E)** Huh7.5.1 cells were co-transfected with pcDNA expression plasmids, GyrB-PKR-WT and the constitutive pCMV-GLuc reporter plasmid, followed by coumermycin A1 treatment as described above. Constitutive luciferase activity was measured as a proxy for protein synthesis and normalized to DMSO-treated controls. Graphs show means +/- standard deviation from at least two independent experiments performed in technical triplicate. Statistical significance was assessed by two-way ANOVA followed by Sidak’s post-hoc test (ns; not significant; ***p<0.001; ****p<0.0001).

We next assessed stress granule (SG) formation by immunostaining for G3BP1. We focused on NS5A from JFH1 and J4 as our models, given that PKR is activated in the context of JFH1 (genotype 2a) infection **(Figure 1A-B)**, while genotype 1b NS5A with J4-like ISDR sequences was previously reported to inhibit PKR. Consistent with the reporter assays, neither JFH1-NS5A nor J4-NS5A reduced SG formation following coumermycin A1 treatment, whereas SPPV-K3L markedly suppressed SG assembly **(Figure 3C-D)**. Lower basal levels of SGs were observed in DMSO-treated cells expressing IAV-NS1 or SPPV-K3L, further supporting the ability of these viral antagonists to suppress PKR activation and downstream SG formation. In contrast, NS5A expression did not alter basal SG levels relative to the GFP control.

We next performed the GyrB-PKR protein synthesis reporter assay in Huh7.5.1 hepatoma cells, which better mimic the natural target cells of HCV infection. However, as observed in the 293T cell model, NS5A failed to rescue protein synthesis following coumermycin-induced Gyr-PKR activation, in contrast to the SPPV-K3L control **(Figure 3E)**. The inability of NS5A to rescue translation or prevent SG formation in this system indicates that HCV NS5A does not inhibit canonical PKR signaling at or downstream of the dimerization step.

### CypA does not regulate downstream PKR activity

The host factor CypA binds to PKR and has been suggested to positively regulate PKR activity [43]. Since CypA also interacts with NS5A [44], and we found that the antiviral potency of Cyp inhibitors partially depended on PKR [33], we next tested the role of CypA in viral PKR antagonism. We generated CypA knockout HEK293T/17 cells using CRISPR/Cas9 **(Figure 4A)**. To investigate the requirement of CypA for PKR downstream signaling, we co-transfected CTRL or CypA KO cells with GyrB-PKR-WT and the constitutive pCMV-GLuc reporter plasmids, together with pcDNA expressing GFP, SPPV K3L or HCV NS5A. Coumermycin-induced PKR dimerization and activation reduced constitutive GLuc activity, reflecting inhibition of protein synthesis, regardless of CypA expression **(Figure 4B)**, indicating that CypA is not required downstream of PKR dimerization. Furthermore, CypA expression did not affect the ability of SSPV-K3L to rescue protein synthesis, and NS5A was unable to rescue protein synthesis in either the presence or absence of CypA.

**Figure 4.**
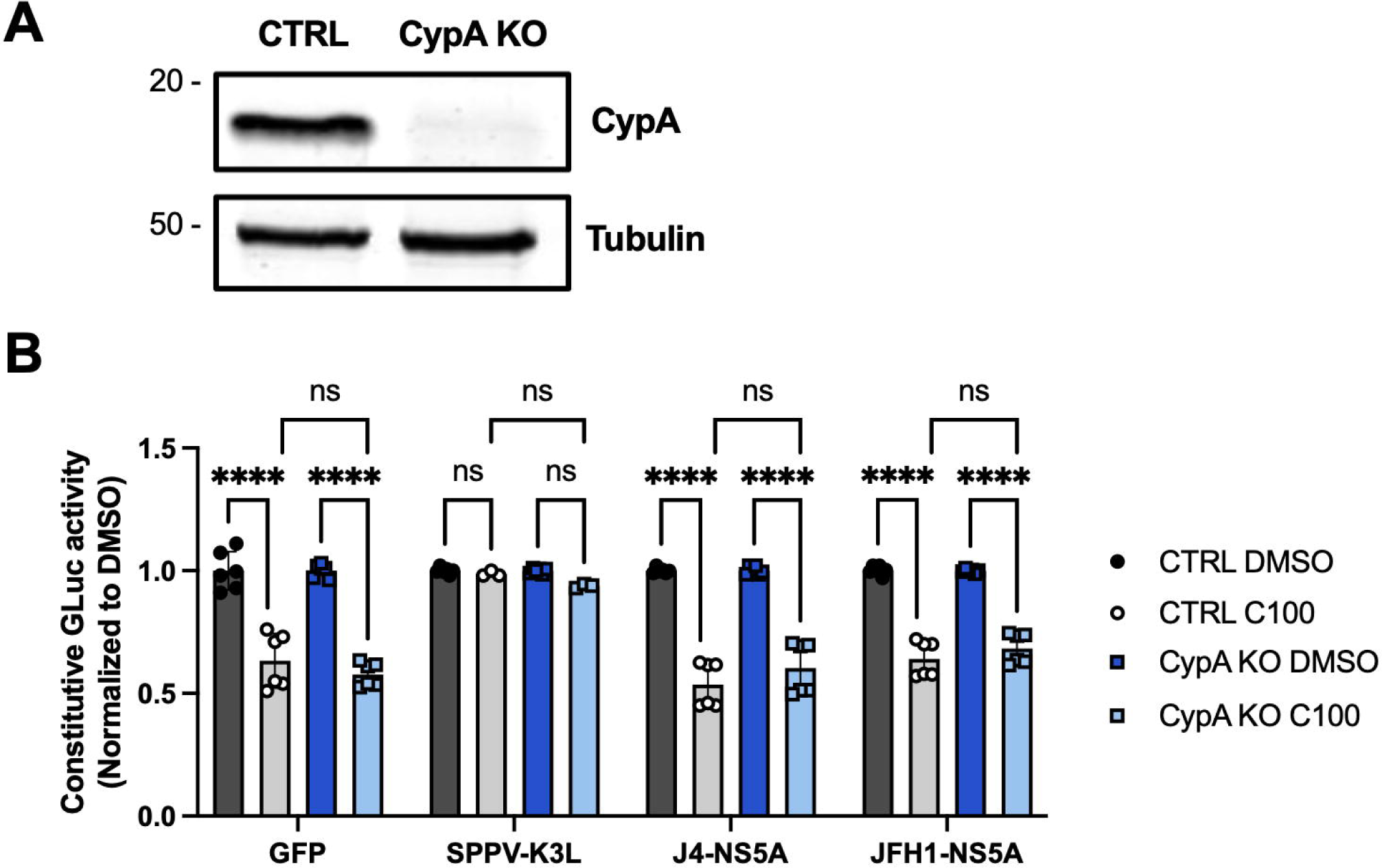
CypA is not required for downstream PKR signaling. **(A)** Knockout of CypA in HEK293T/17 cells was confirmed by western blot. **(B)** HEK293T/17 CTRL or CypA-KO cells were co-transfected with 500 ng pcDNA expression plasmids encoding GFP, SPPV-K3L, JFH1-NS5A-FLAG or J4-NS5A-FLAG, together with 500 ng of GyrB-PKR-WT and the pCMV-GLuc reporter plasmid. The next day, cells were treated with DMSO vehicle or coumermycin A1 (100 ng/mL) for an additional 24 h and luciferase activity was measured and normalized to DMSO-treated control.

### NS5A does not inhibit dsRNA-induced PKR activation

To determine whether NS5A inhibits PKR activation at an upstream step by interfering with dsRNA binding to PKR, HEK293T/17 cells were co-transfected with expression plasmids encoding GFP, JFH1-NS5A, J4-NS5A, SPPV-K3L, or IAV-NS1. The next day, cells were transfected with the synthetic dsRNA analog poly(I:C) for 6 h to induce PKR activation. Notably, both IAV-NS1 and SPPV-K3L reduced basal PKR activation in unstimulated cells, consistent with their established roles as PKR antagonists, whereas NS5A had no effect on PKR phosphorylation **(Figure 5A)**. Given the relatively low transfection efficiency in these experiments, we could not reliably assess the impact of the viral proteins on global poly(I:C)-induced PKR activation by western blot **(Figure 5A)**. However, neither NS5A construct rescued PKR-dependent translational shutdown, as measured by stress granule formation **(Figure 5B-C)**, in contrast to IAV-NS1 and SPPV-K3L, both of which inhibited poly(I:C)-induced SG assembly. Basal SG levels were also reduced in cells expressing IAV-NS1 or SPPV-K3L, further supporting their ability to suppress basal PKR activation.

**Figure 5.**
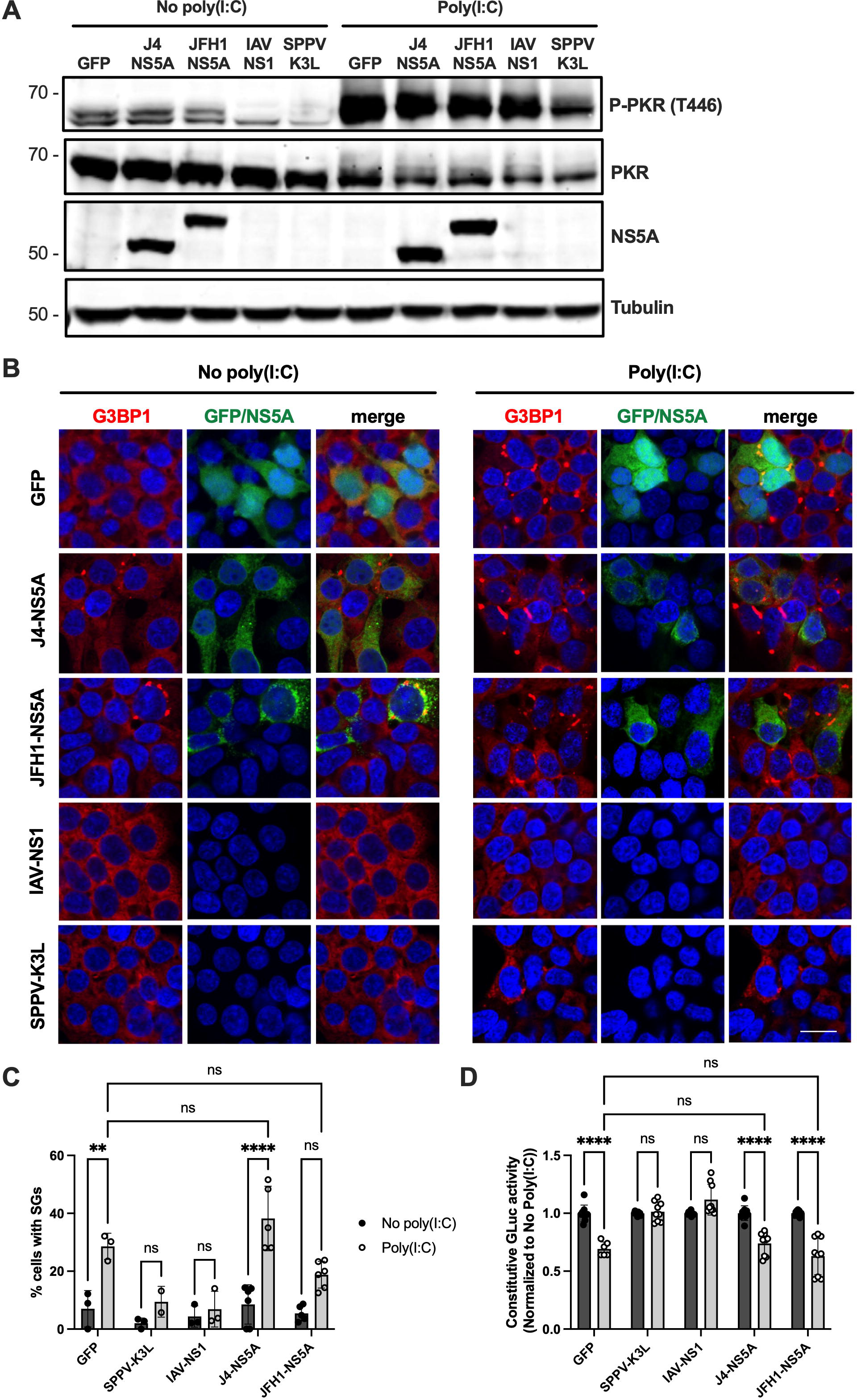
NS5A does not inhibit dsRNA-induced PKR activation. **(A)** HEK293T/17 cells were co-transfected with 500 ng pcDNA expression plasmids encoding GFP, JFH1-NS5A-FLAG, J4-NS5A-FLAG, SPPV-K3L or IAV-NS1. After 24 h, cells were transfected with 2 μg of poly(I:C) for 6 h. Cells were then processed for western blot to detect PKR activation, using antibodies against phosphorylated PKR T446, total PKR, NS5A and tubulin. **(B-C)** SG formation of transfected cells described above were observed by immunofluorescence following staining for G3BP1 (red) and NS5A (green) **(B)**. **(C)** The percentage of cells with SGs was quantified in at least three independent fields of view. **(D)** HEK293T/17 cells were transfected as described above along with the pCMV-Gluc reporter plasmid. Following poly(I:C) stimulation, constitutive luciferase activity was measured as a proxy for protein synthesis.

To isolate the effect of poly(I:C) stimulation to transfected cells expressing the viral proteins, we co-transfected cells with the pcDNA expression vectors encoding GFP, SPPV-K3L, IAV-NS1, J4-NS5A or JFH1-NS5A together with the constitutive pCMV-GLuc reporter plasmid. In control cells, poly(I:C) transfection resulted in decrease in constitutive luciferase activity, consistent with PKR-mediated translational halt. The inhibitory effect of poly(I:C) on protein synthesis was rescued in cells expressing SPPV-KL3 or IAV-NS1, but not NS5A **(Figure 5D)**. Together, these findings indicate that HCV NS5A does not inhibit PKR activation by preventing dsRNA-mediated activation of PKR and further support the conclusion that NS5A does not directly antagonize PKR by blocking dsRNA binding or inhibiting PKR dimerization.

### NS5A does not inhibit PKR activation in the presence of other non-structural proteins

During infection, NS5A functions within the viral replication complex and interacts with other HCV non-structural (NS) proteins. Therefore, we next investigated whether the presence of additional viral NS proteins could influence the ability of NS5A to regulate PKR activation. Huh7-Lunet/T7 cells were transfected with chimeric pTM constructs encoding HCV non-structural proteins of different genotypes, from NS3 to the 3′ end. We tested pTM vectors expressing NS3-5B from genotype 1b (GLT1 and Con1), genotype 2a (J6 and JFH1) or genotype 3a (S52) strains. No additional PKR activator was added, since expression of NS5B has been shown activate PKR through dsRNA generated by its RNA-dependent RNA polymerase activity [40]. Furthermore, transfection of plasmid DNA can activate PKR, which has been attributed to production of dsRNA formed by positive-sense and negative-sense mRNA transcription from cryptic promoters [45–47], enabling us to assess PKR activation only in pTM transfected cells expressing NS3-5B or GFP control. At 24 h post-transfection, PKR activation was assessed by western blot. Regardless of the genotype, PKR activation was not reduced in the presence of NS5A or other HCV non-structural proteins **(Figure 6A)**. Consistent with these findings, confocal microscopy analysis of the SG marker eIF3η revealed SG formation in NS5A-expressing cells **(Figure 6B)**. Together, these results indicate that NS5A, regardless of genotype, does not inhibit PKR activation or SG formation, even in the context of other viral non-structural proteins.

**Figure 6.**
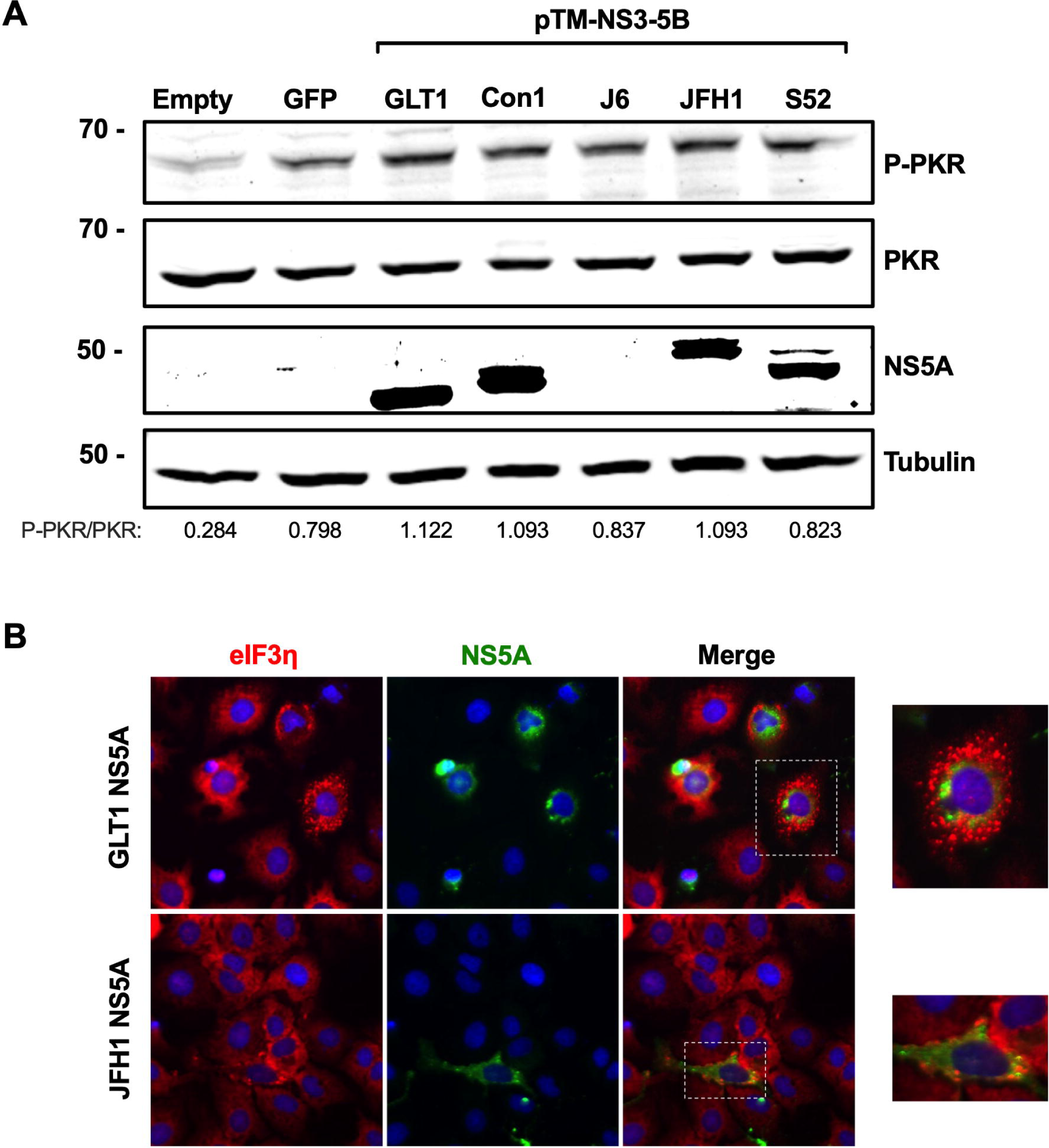
NS5A does not inhibit PKR in the presence of other HCV non-structural proteins. **(A)** Huh7-Lunet/T7 cells were transfected with 500 ng of the pTM constructs encoding NS3-5B from the indicated genotypes. After 24 hours, PKR activation was assessed by western blot for phosphorylated PKR, total PKR, NS5A and tubulin. Band density analysis was performed to determine the ratio of P-PKR to total PKR. **(B)** SG formation was visualized by confocal microscopy for the SG component eIF3η (red), NS5A (green), with DAPI nuclear stain

## DISCUSSION

Responsiveness to IFN-based therapy varies across HCV genotypes [48], and early evidence suggested a role for the NS5A-PKR interaction in determining IFN sensitivity. Gale et al. showed that NS5A from IFN-resistant genotype 1b strains can directly bind to the catalytic domain of PKR independently of dsRNA and disrupt its dimerization, thereby preventing downstream eIF2α phosphorylation and translational arrest [21, 22, 49]. However, PKR is activated by HCV JFH1 infection [25] or RNA replication [50], despite robust NS5A expression, arguing against NS5A as an effective PKR inhibitor. Indeed, PKR activation has been shown to promote HCV infection through SG formation [26, 27] and halting translation of IFNs and ISGs [29]. To reconcile these observations, we sought to investigate the potential genotype dependence and mechanism of NS5A-mediated PKR antagonism. We systematically examined the effect of NS5A on PKR activation using orthogonal assays in cell culture models and found little evidence that NS5A functions as a direct inhibitor of PKR-induced translational halt.

To directly address whether NS5A inhibits PKR activation, we employed a chemically inducible GyrB-PKR dimerization system that bypasses dsRNA sensing and selectively activates PKR through forced dimerization [36]. This system effectively recapitulates canonical PKR signaling, including eIF2α phosphorylation, translational repression, and SG formation **(Figure 2)**. Using this approach, we found that NS5A from multiple HCV strains and genotypes failed to rescue protein translation or prevent SG formation following PKR activation **(Figure 3)**. In contrast, the poxvirus PKR antagonist SPPV-K3L strongly inhibited PKR-eIF2α signaling, as expected [42]. We further tested whether NS5A might inhibit PKR upstream of dimerization by interfering with dsRNA sensing. However, neither JFH1-NS5A nor J4-NS5A reduced PKR activation, protein translational shutdown or SG formation, while IAV-NS1 and SPPV-K3L suppressed PKR activation **(Figure 5)** as expected [18, 42]. These findings argue against a model in which NS5A antagonises PKR by interfering with dsRNA binding or PKR dimerization, which are typical mechanisms used by other viral PKR antagonists.

Our findings differ from previous observations that NS5A inhibits PKR dimerization [21, 22], which could be explained by different experimental approaches and model systems. While these earlier studies were instrumental in identifying a possible interaction between PKR and NS5A, the use of *in vitro* assays with recombinant proteins, coupled with yeast-based functional assays, may not fully recapitulate the physiological cellular context of HCV infection. While an interaction between NS5A and PKR in Cos-1 African green monkey kidney cells was also identified [22], the use of an N-terminal FLAG tag in the NS5A expression construct in this model may have disrupted NS5A localization to endoplasmic reticulum (ER) membranes, which is mediated by its N-terminal amphipathic helix [51, 52]. In contrast, we used C-terminally FLAG-tagged NS5A, as modifications to the C-terminus are not expected to affect ER localization [52]. We speculate that the localization of NS5A to ER membranes and within the replication organelle may normally limit its ability to interact with cytoplasmic PKR.

He et al. subsequently used recombinant vaccinia virus (VACV) expressing NS5A in place of the poxvirus E3L PKR antagonist to investigate the role of NS5A in PKR inhibition following pre-treatment with exogenous IFN [49]. In this model, NS5A expression transiently inhibited PKR activation and downstream translational arrest, but the inhibitory effect was lost by 6 hpi infection in VACV-NS5A-infected cells, compared to 4 hpi in the VACV-ΔE3L control. The transient nature of the observed inhibition suggests that NS5A does not effectively antagonise PKR-mediated translational halt in the context of a DNA virus infection that generates dsRNA from viral gene transcripts [53].

Evidence that PKR-mediated translational halt is antagonised by NS5A during authentic HCV infection is still lacking. To address this question, we initially attempted to use NS5A chimeric viruses based on the J6/JFH1 backbone [38]. However, differences in replication efficiency and kinetics among these chimeric viruses precluded the ability to accurately compare PKR activation or antagonism (data not shown). Instead, we used a polyprotein expression system to evaluate the impact of NS5A from different genotypes on PKR activation in the context of the other viral non-structural proteins that form the HCV replicase complex, including NS5B, which has been shown to activate PKR through its RNA-dependent RNA polymerase activity [40]. Even when expressed in the more authentic context of NS3-5B, NS5A did not suppress PKR activation or SG formation, regardless of genotype or strain **(Figure 6)**.

Our finding that NS5A does not inhibit dsRNA-mediated PKR activation is consistent with previous studies showing that NS5A preferentially binds single-stranded viral RNA rather than double-stranded RNA replication intermediates [54]. Therefore, NS5A is not expected to directly sequester dsRNA. Furthermore, HCV RNA replication occurs within membrane-associated replication organelles that compartmentalize viral RNA and limit its accessibility to cytoplasmic RNA sensors [55]. However, the highly structured IRES in the 5′ UTR from HCV genotype 1b has been shown to activate PKR in vitro [56], indicating that RNA secondary structures in the viral genome could contribute to PKR activation in infected cells. In the same study, NS5A binding to the HCV IRES decreased activation of PKR, as measured by in vitro activation assays. Interestingly, Karamichali et al. demonstrated that NS5A down-regulates HCV IRES activity in HepG2 cells, but not Huh7 cells, which was attributed to PKR inhibition [57]. The functional relevance of these findings remains to be demonstrated in the context of HCV infection.

While our study focused on PKR-eIF2α-mediated translational repression, PKR also regulates antiviral transcriptional responses driven by NF-κB and IRF1 [58]. Zhang et al. demonstrated that silencing PKR enhances HCV genotype 1a replication by inhibiting NF-κB activation and therefore attenuating the type I IFN response [59]. HCV NS5A was also shown to block the activation of IRF1 and inhibit IRF1-dependent gene expression [14]. It was recently shown that NS5A domain I modulates the PKR-IRF1 signaling axis to counteract a block to viral assembly [60]. Our finding that cyclophilin inhibitors appear to enhance IRF1-dependent antiviral gene expression in HCV-infected cells in a PKR- and IRF1-dependent manner [33] suggests a potential role for the CypA-NS5A or CypA-PKR interaction in regulating this pathway. While the specific mechanisms through which PKR activates or regulates these downstream signalling pathways remain to be determined, NF-κB activation by PKR requires neither its kinase activity nor the PKR-eIF2α axis [17]. Another study found that PKR associates with the mitochondria antiviral signaling protein (MAVS) and the TRAF3 adaptor protein early during HCV infection to promote induction of IRF3-dependent genes such as ISG15 [61]. This interaction required the double-stranded RNA binding region of PKR, but not its catalytic activity. Overall, further studies are needed to elucidate how NS5A modulates these signaling pathways downstream of PKR, which will also inform our understanding of the role of PKR as a signalling adaptor protein.

Accumulation of mutations in the recently identified replication enhancing domain (ReED) was shown to confer high replication fitness to a genotype 1b clinical HCV isolate, GLT1 [24]. Since the ReED maps to a region of NS5A that contains the ISDR and CypA binding site, as well as the proposed PKR binding domain, we evaluated whether NS5A from GLT1 was able to inhibit the PKR-eIF2α pathway more effectively than other strains. However, despite the mutations in the ISDR, GLT1 NS5A did not inhibit PKR phosphorylation, downstream translational halt, or SG formation **(Figure 3A, 6A-B)**. Historically, mutations in the ISDR region conferred sensitivity or resistance to IFN therapy, which was attributed in part to PKR inhibition. However, we speculate that modulation of replication fitness, rather than a specific effect on the proposed NS5A-PKR interaction, likely underlies the role of ISDR mutations in IFN sensitivity. A recent study found that high replication fitness may contribute to, but likely does not fully explain, the response to IFN treatment [62].

Overall, we found that NS5A, regardless of the tested genotype or strain, has limited ability to antagonize the canonical PKR-eIF2α pathway. While earlier studies provided evidence supporting an interaction between NS5A and PKR, our data indicate that NS5A does not effectively inhibit PKR activation and SG formation during HCV JFH1 infection and in cell culture models with ectopic expression of NS5A from different strains. Future studies using replication-competent models and approaches that incorporate authentic HCV RNA structures will help to further clarify potential contributions of NS5A to PKR regulation in specific contexts during HCV infection. Additionally, further studies are needed to understand the potential roles of NS5A in regulating non-canonical PKR signaling pathways. Collectively, these findings refine our understanding of HCV evasion mechanisms by demonstrating that NS5A is not a robust antagonist of classical PKR-eIF2α signaling.

## ACKNOWLEDGEMENTS

This work was supported by grants to CCC from the Natural Sciences and Engineering Research Council of Canada (RGPIN-2020–04277) and the Canadian Institutes of Health Research (PTT-184008). We are grateful to the Canadian Foundation for Innovation John R. Evans Leaders Fund (CCC) for equipment that supported this research. VL acknowledges funding from the Deutsche Forschungsgemeinschaft (DFG), grant number 272983813. CEGF acknowledges a MITACS Globalink Graduate Fellowship and a PhD fellowship from the Canadian Network on Hepatitis C (CanHepC), as well as the CanHepC-TRR179 student internship exchange program. CanHepC is funded by a joint initiative of the Canadian Institutes of Health Research (NHC-142832, NHE-174228, HPC-178912) and the Public Health Agency of Canada. ERK acknowledges a Canada Graduate Scholarship (Master’s Program) from the Canadian Institutes of Health Research. We are grateful to Dr. Alessia Ruggieri (Heidelberg University) for helpful discussions. We thank Dr. Craig McCormick (Dalhousie University), Dr. Thomas Dever (National Institutes of Health) and Dr. Judith Gottwein (University of Copenhagen) for kindly sharing plasmids that supported this work.

